# Visualization of the Dynamics of Histone Modifications and Their Crosstalk Using *PTM-CrossTalkMapper*

**DOI:** 10.1101/801837

**Authors:** Rebecca Kirsch, Ole N. Jensen, Veit Schwämmle

## Abstract

Visualization of large-scale, multi-dimensional omics data is a major challenge. Experimental study designs in proteomics research produce multiple data layers, e.g. relationships between hundreds of proteins, their interactions and abundances as a function of time or perturbation, as well as dynamics of post-translational modifications (PTMs) and allelic variants (proteoforms). These different levels and types of information of proteins and proteoforms complicate data analysis and generation of insightful and comprehensible graphic visualizations. Middle-down mass spectrometry of histone proteins now allows quantifying hundreds of histone proteoforms including co-existing methylation, acetylation and phosphorylation events at distinct amino acid residues within histone molecules. The histone PTM landscape plays a dominant role in the regulation of chromatin activity and transcriptional and epigenetic control. The dynamics of these reversible modifications are governed by reader, writer and eraser enzymes that cooperate to regulate molecular mechanism that rely on multiple interdependent PTM marks in histones and nucleosomes in chromatin. This PTM crosstalk can be quantified and provides a detailed picture of the underlying rules for setting the histone PTM landscape and chromatin activity, and is available to the community via our CrosstalkDB platform. Here, we developed a new computational method, *PTM-CrossTalkMapper*, to visualize the dynamics of histone PTMs for different experimental conditions, replicates and proteoforms onto a landscape, thereby describing the crosstalk and interplay between PTMs in a more comprehensible manner. We show the power of combining different levels of information on such crosstalk maps for histone PTM dynamics in mouse organs during the aging process. The *PTM-CrossTalkMapper* toolkit provides flexible functions to create these maps in various scenarios of multi-dimensional experimental designs, including histone PTM patterns and PTM crosstalk.

The source code is available at https://github.com/veitveit/CrossTalkMapper.

## Introduction

With the development of increasingly efficient mass spectrometry technologies, proteomics experiment design involve multiple levels of experimental and biological factors. For example, the proteomes of samples derived from different tissues - in technical and biological replicates - can be measured at different conditions and at multiple time points. Large-scale proteomics studies thereby deliver deep and rich information about the investigated system but create new challenges in data analysis and reporting. Generating biological insight is further complicated when studying post-translational modifications (PTMs) where each individual protein within a sample often comprise several proteoforms that carry multiple, different PTMs that vary in their relative abundances, that in turn regulate or fine-tune protein function. This additional level of data complexity introduced by PTMs is particularly pronounced for heavily modified proteins like histones. Combinatorial explosion leads to a large variety of different histone proteoforms of which many are known to have mutually dependent biological functions. The development of tools to visualize these complex, multi-layered data has been lagging behind and thus prevents discovery and insightful presentation of inherent biological information. There is a need for novel approaches of PTM data representation [1,2], which we are addressing in this manuscript. Histone proteins carry complex epigenetic information in their many differently modified proteoforms. Histones are also involved in basic cellular processes, including chromatin structure and function and transcriptional regulation of gene expression. This control is achieved by enzymatic PTM readers that recognize specific PTMs and recruit PTM writers and erasers to respectively add or remove modifications at other sites based on the already present PTMs [3,4]. This coordination and cooperation between multiple PTMs in histones (cis or trans) is referred to as PTM crosstalk, which can be positive or negative depending on how the PTMs influence each other (see Fig. 1A). One well-established example of PTM crosstalk is phosphorylated serine 10 on histone H3 (H3S10p, which is bound by the histone acetyltransferase Gcn5, leading to the acetylation of lysine 4 on H3 (H3K4ac) leading to destabilization of the nucleosome and upregulation of the associated promoter [5,6]. Directly measuring the crosstalk that is taking place between PTMs under certain circumstances is a major challenge, as it requires precise information of PTM occupancy at multiple, distinct amino acid residues on a histone, i.e. PTM co-occurrence. Traditional bottom-up mass spectrometry of small peptides of 8-20 amino acid residues cannot reliably retain this co-occurrence information, while the top-down approach applied to entire histones is not sensitive enough to distinguish specific PTM combinations from each other due to sample complexity. Middle-down mass spectrometry (see Fig. 1B) became established to measure the set of histone tail proteoforms present in a sample and the relative proteoform abundances can be determined [2,7,8], including also mammalian organisms and tissues [9]. Based on these histone proteoform profiles, the abundances of individual and combinatorial PTMs in the sample can be calculated. The crosstalk of a combination of two PTMs is then defined as positive if both are observed together on the same histone more often than expected by chance, and negative if they are detected together more rarely than expected by chance. We define the observed vs. expected co-occurrence ratio as the interplay score and used it to assess the type and strength of PTM crosstalk [10].

**Figure 1:**
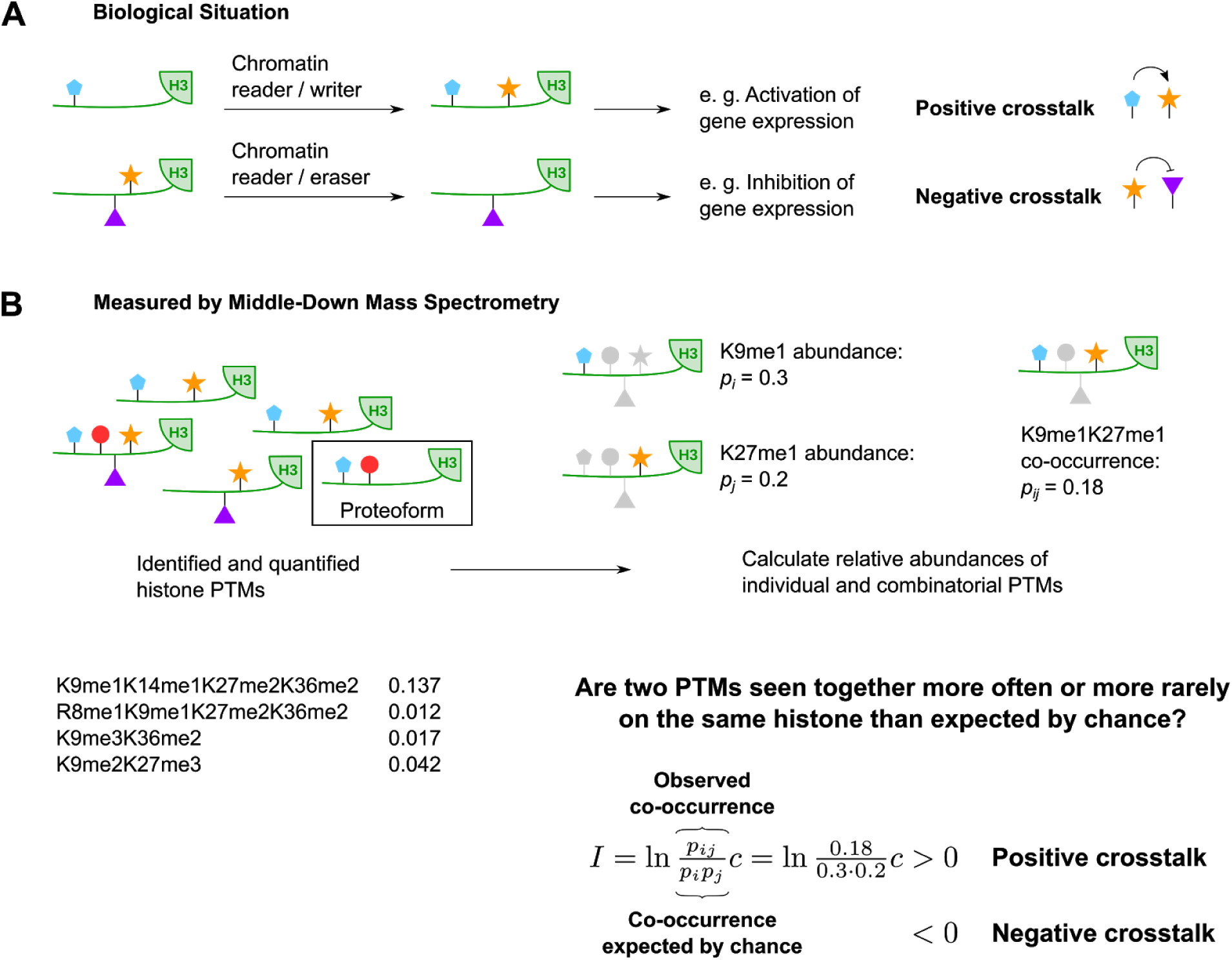
Crosstalk concept and definition of interplay score. A) Chromatin writers and erasers place and remove histone PTMs based on the recognition of other PTMs by chromatin readers. This (non-physical) interaction between PTMs is referred to as positive or negative PTM crosstalk, depending on how the PTMs influence each other. Histone PTM crosstalk dynamics mediate the expression of histone-associated genes. B) The information depth of middle-down mass spectrometry enables measuring PTM crosstalk using the previously introduced interplay score [10]. I: interplay score, c: correction factor.

The interplay score has been utilized in the analysis platform CrosstalkDB (http://crosstalkdb.bmb.sdu.dk) [10,11], which hosts a collection of quantitative histone PTM datasets and allows uploading mass spectrometry datasets containing quantified proteoforms or long modified peptides. It provides a number of statistical analyses of PTM distributions and crosstalk for each dataset or dataset combination of interest.

For a comprehensive understanding of PTM functions, however, comparing the dynamics of multiple PTM combinations and crosstalk under different conditions is crucial. At the same time, it requires keeping track of individual and combinatorial PTM abundances. Previous approaches to visualize combinatorial PTM data [9–13] are able to provide an overview over the set of identified PTM pairs and an additional parameter, e.g. their crosstalk, but only for one condition at a time or an average over conditions.

To boost the efficient comprehension of complex, multi-layered PTM datasets, we present a novel visualization framework, *CrossTalkMapper* (https://github.com/veitveit/CrossTalkMapper), which serves as an extension to CrosstalkDB. We adjusted the interplay score to provide more accurate quantitative crosstalk values, making it possible to generate a symmetric crosstalk landscape onto which multiple PTM pairs can be projected with their crosstalk and abundances. Data transformation allows combining the differing PTM landscapes for several conditions into one crosstalk map. This flexible visualization tool allows the comparison of temporal PTM dynamics for multiple PTM pairs between different tissues or cell types and for various perturbations. This simplifies the identification of interesting combinatorial PTM features and differences between conditions, providing direction and insights for disentangling the underlying epigenetic mechanisms.

## Methods

### Dataset and data preparation

Relative proteoform abundances of histone H3 tails from brain, heart, kidney and liver of 3, 5, 10 and 18 months old mice had previously been determined for histone variants H3.1/2 and H3.3 by middle-down mass spectrometry [9] and were downloaded from CrosstalkDB (http://crosstalkdb.bmb.sdu.dk) [10,11]. Columns for tissue and replicate number were added to the file based on the dataset name and ID. 0-imputation of missing values was performed for biological replicates (4 for each of the 3, 5 and 10 month samples, 2 for each of the 18 month samples). For plots comparing histone variants, proteoform relative abundances were normalized with respect to the total abundance of the respective variant. Otherwise, proteoform abundances for the individual variants were summed to obtain H3 total proteoform abundances. Then, biological replicates were averaged unless stated otherwise.

### Abundance-corrected interplay score

The interplay score as a measure for crosstalk between two PTMs *i* and *j* has previously [10] been defined as

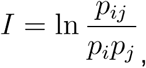

where *p*_*i*_ and *p*_*j*_ are the total abundances of PTMs *i* and *j*, respectively, relative to the number of all histone tails present in the sample. These individual abundances are calculated by summing the relative abundances for all proteoforms containing the respective PTM, e.g K9me1 or K27me1. The product of the individual abundances is equal to the random chance of both modifications being observed on the same histone tail.*p*_*ij*_ is the relative observed co-occurrence of both modifications on the same histone tail, obtained by summing all relative abundances of proteoforms containing both modifications, e.g. K9me1K27me1. Positive crosstalk between PTMs *p*_*i*_ and *p*_*j*_ is assumed for *I > 0*, and negative if the interplay score is < 0.

To represent positive as well as negative crosstalk over the whole range of possible abundance values more accurately, the interplay score was adjusted as

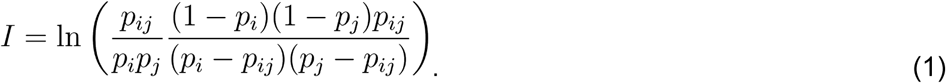

While the principle and the interpretation of the final score remain as for the previous version, the corrected score ensures symmetry and correct calculation of the limiting cases describing maximal positive or negative crosstalk.

### New multi-layer visualization approach

Individual PTM abundances *p*_*i*_ and *p*_*j*_ are transformed into 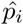 and 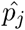 as

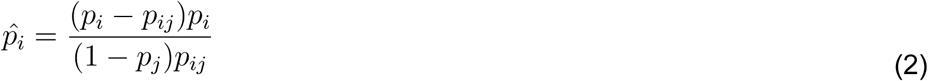

and

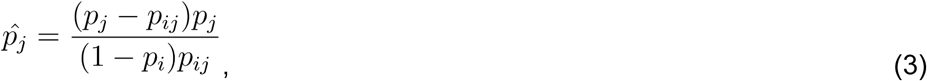

where *p*_*i*_, *p*_*j*_, *p*_*ij*_ ≠ 0 Using **geom_raster**() from the **ggplot**() package, a log-log scale raster plot is generated with 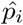 and 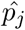 coordinates ranging from the minimum to maximum transformed abundance contained in the dataset and the respective interplay score, which is given by

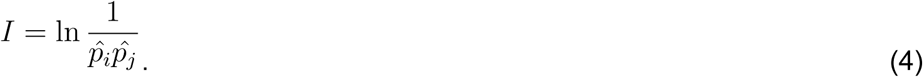

By having only two variables for the crosstalk, it can be projected onto an invariant crosstalk landscape. Transformed data points from multiple PTM pairs and different samples, e.g. tissues, time points or replicates, can be projected onto the same crosstalk map. Data point colors can be used to encode abundances or conditions in a flexible way.

### Open source toolkit

Three main R functions have been implemented in *PTM-CrossTalkMapper* that simplify the workflow from data preparation to plot generation. **prepPTMdata**() handles data that can be imported from CrosstalkDB or from a file that contains proteoform abundances. The data is then prepared for further processing. This includes 0-imputation of missing values and options for normalizing proteoform abundances with respect to histone variants and averaging replicates. Using **calcPTMab**(), non-transformed and transformed individual abundances, co-occurrences and interplay scores are calculated for each observed PTM pair. The function output is a data table containing all pairwise PTM combinations and their parameters, from which data points of interest are selected by the user (an example is provided together with the *PTM-CrossTalkMapper* code) and passed to **CrossTalkMap**(). This function has several options to define how the different data parameters will be encoded in the plot, which sets of data points should be connected and based on which parameter the plot should be divided into subplots, if applicable.

In addition, **line_ab**() and **line_ct**() provide more detailed views of abundances and interplay scores for specific PTMs and PTM pairs.

## Results

### Refined definition of the interplay score

We introduced the interplay score in 2014 [10] to assist in the identification of co-occurring PTMs based on accurate quantitative mass spectrometry experiments. Here, we refine the interplay score in order to provide a symmetric model. The new definition of the interplay score avoids underestimation of positive crosstalk by too low interplay scores when there is a high co-occurrence of PTMs relative to their individual abundance, i.e. a large fraction of the histones carrying one PTM are also decorated by the second PTM (see Fig. 2). By improving this symmetry, crosstalk measures for lowly and highly abundant PTMs become more comparable. This was achieved by adding a correction factor which will also be used to build the crosstalk maps we introduce below. This leads to a better visual representation of PTM changes with respect to their crosstalk.

**Figure 2:**
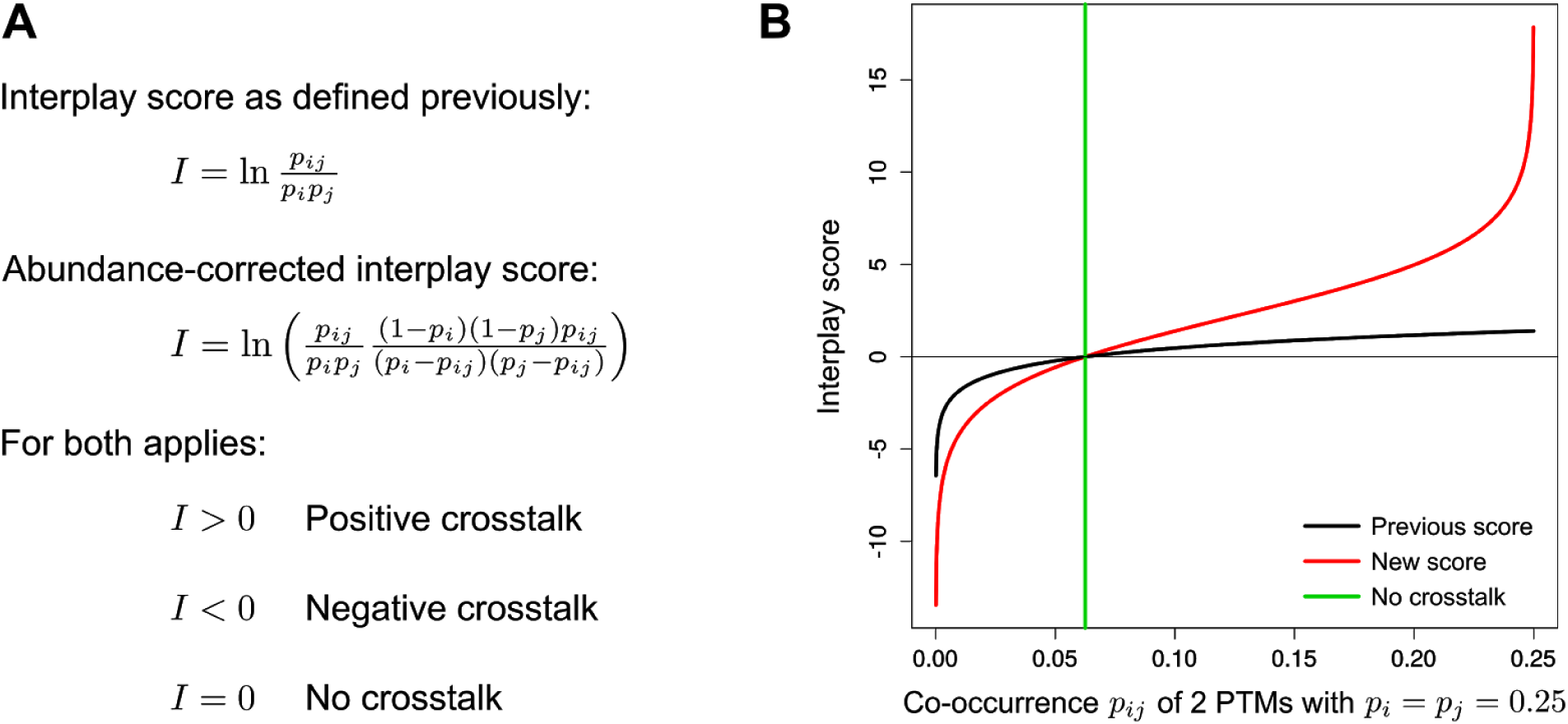
Revised interplay score definition. We have adapted the previously defined interplay score to better represent extreme crosstalk cases where almost all histones carrying one PTM are decorated by the second PTM as well.

With the revised score version, extreme cases are now described accurately. For instance, mutual exclusion of PTMs (e.g. acetylation and methylation on the same residue) corresponds to an interplay score of minus infinity. Such negative crosstalk is an intrinsic (chemical) property of the system that cannot be quantified from mass spectrometry data. Maximal positive crosstalk now also is described by an extreme interplay score of infinity. In this scenario, a specific PTM cannot be found without another PTM, corresponding to an extreme case where i) histones are only modified with a specific PTM if the histone is already modified with the crosstalking PTM and ii) removal of the PTM also leads to simultaneous removal of the other PTM. We consider this case to be rare.

Hence, the interplay score restricts to quantifiable and realistic PTM crosstalk in a symmetric manner and lays the foundation for the crosstalk maps introduced in this manuscript.

### Definition of a PTM crosstalk map

The adapted interplay score makes it possible to create a symmetric crosstalk landscape that is defined by the abundances of the PTMs contained in a dataset. Data points corresponding to any PTM pair on the same protein can be projected onto this landscape, and thus create what we refer to as a *crosstalk map*.

To simultaneously plot arbitrary combinations of PTMs and conditions, PTM abundances are transformed. This allows overlaying PTM patterns from different conditions such as tissues or time points. On the resulting crosstalk map, the crosstalk dynamics of multiple PTM pairs can easily be followed over time, across tissues or within replicates.

Fig. 3A depicts the abundances of mono-methylated Lys 9 and Lys 27 of histone H3 (H3K9me1, H3K27me1) from mouse heart tissue and their temporal change across four time points (3, 5, 10 and 18 months). This simple line plot shows the co-occurrence of both PTMs as well as the corresponding interplay score according to Eq. (1). The fraction of histones carrying only one of these marks decreases with age while histones decorated with both PTMs become slightly more abundant. This rather drastic change in the PTM composition correlates with a strong increase of the interplay score which changes from negative crosstalk, where the PTMs are mutually exclusive, to their independent setting, given by an interplay score of 0. While Fig. 3A is quite intuitive, addition of other PTM pairs and additional experimental conditions will require adding new figures and thus make direct comparison of PTM frequency and their crosstalk over different scenarios increasingly difficult.

**Figure 3:**
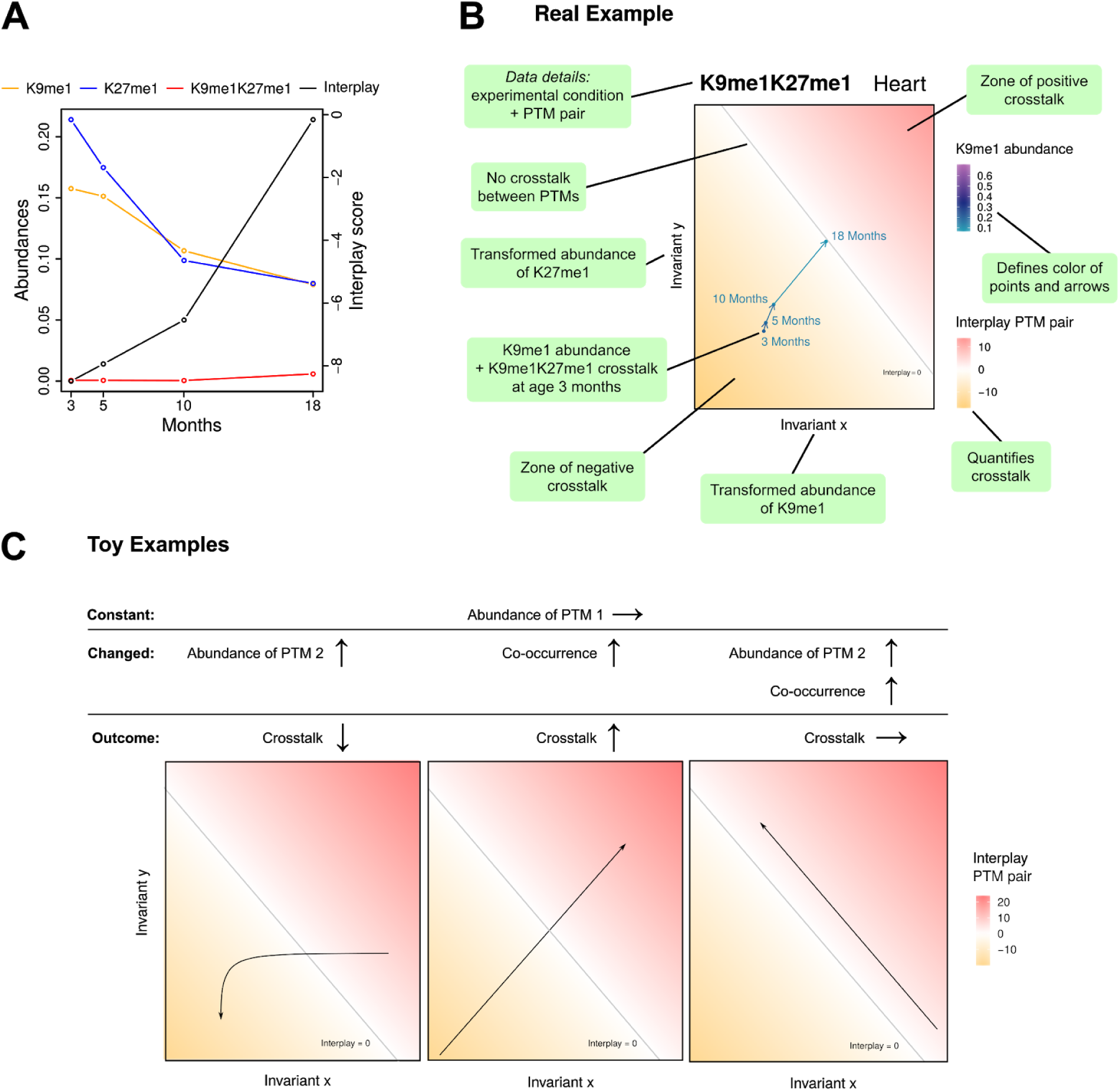
Concept of PTM crosstalk maps. A) The change of individual and combinatorial abundances as well as the resulting crosstalk of one specific PTM pair over time at one condition (heart). B) A simple example of a crosstalk map showing the same data, together with annotations of all plot elements. In this example, K9me1 abundance is color-coded but the tool allows to flexibly encode either of the individual and combinatorial abundances like this. C) Influence of individual crosstalk parameters on the data point trajectory. Artificial toy examples assume the abundance of one PTM to be constant, while either the abundance of the second PTM, the co-occurrence or both parameters are changed. In this ideal setting, the trajectories and changes in crosstalk observed in the crosstalk map would look as indicated.

PTM crosstalk maps allow such an overall comparison by providing a common visualization scheme where multiple PTMs and different experimental conditions are projected onto the same crosstalk landscape. This is illustrated in Fig. 3B, where the dynamics of PTM abundance and their crosstalk from the example in Fig. 3A can be recognized by their trajectory on the crosstalk landscape. At the same time, the crosstalk landscape leaves room for additional PTM pairs and conditions as shown below. We consider this new visualization a practical method to compare PTM dynamics across nearly arbitrary experimental conditions and PTM combinations.

PTM crosstalk maps illustrate changes in the histone landscape by trajectories of combinatorial PTMs. Fig. 3C shows artificial “toy examples” with controlled settings of the crosstalk parameters. We assumed the abundance of one PTM to be fixed and changed either the second PTM individual abundance, co-occurrence or both, to observe the resulting crosstalk change and behavior in the crosstalk landscape. Increasing co-occurrence exhibits a similar behavior when compared to the example shown in Fig. 3B, showing a linear trajectory perpendicular to the diagonal indicating absence of PTM crosstalk. Biologically, this could correspond to situations where two PTMs interact indirectly through the enzymes responsible for their deposition or removal: i) histones with one of the PTMs recruit writer molecules to add the second PTM, e.g. by an increased enzyme affinity; ii) histones simultaneously modified by both PTMs become less accessible to the enzymes that remove these PTMs. Hence, the trajectories can be interpreted by corresponding specific biological control processes that dominate the histone PTM landscape.

### PTM-CrossTalkMapper: Visualization of multi-dimensional PTM datasets

The multi-layered visualization approach has been implemented in *PTM-CrossTalkMapper*, a flexible open source toolkit of functions in the R software environment. It contains three main functions that simplify the workflow from data preparation to plot generation, and additional functions to display selected changes of PTM abundances and their crosstalk.

The tool can import PTM datasets stored in the CrosstalkDB database or csv files containing the relevant fields for the relative abundance profiles of the different proteoforms. The first two functions, **prepPTMdata**() and **calcPTMab**(), facilitate data preparation and the calculation of individual and pairwise PTM abundances, while the third main function produces crosstalk maps containing all PTM pairs of interest. The user can select the layers of information that should be displayed and also specify details how they will be encoded. To avoid figures with too dense content, a map can be split into several plots. This highly configurable layout allows to, for example, create separate crosstalk maps per tissue, separate maps for different time points, or comparison across histone isoforms such as H3.1 and H3.3. Further details like PTM abundances or conditions can be displayed by color codes and manual labelling. Trajectories of the combinatorial marks are shown by arrows or lines. Additional R functions provide the option for more detailed views of the parameters of specific PTM pairs changing over time in additional plots (for example, see Fig. 3A).

The R language provides a flexible framework for the selection of PTMs and automation of plot generation for larger datasets. *PTM-CrossTalkMapper* is available on GitHub (https://github.com/veitveit/CrossTalkMapper), along with documentation, auxiliary scripts and code examples of some common use cases that also should help inexperienced R users to get started.

### Application of PTM-CrossTalkMapper to studies of dynamic PTMs in aging research

To demonstrate our PTM visualization framework, we applied *PTM-CrossTalkMapper* to a previously published mass spectrometry dataset, which was derived from four different mouse tissues at four time points (see Fig. 4) to study how aging affects the histone PTM landscape and thus epigenetic control of major biological processes [9].

**Figure 4:**
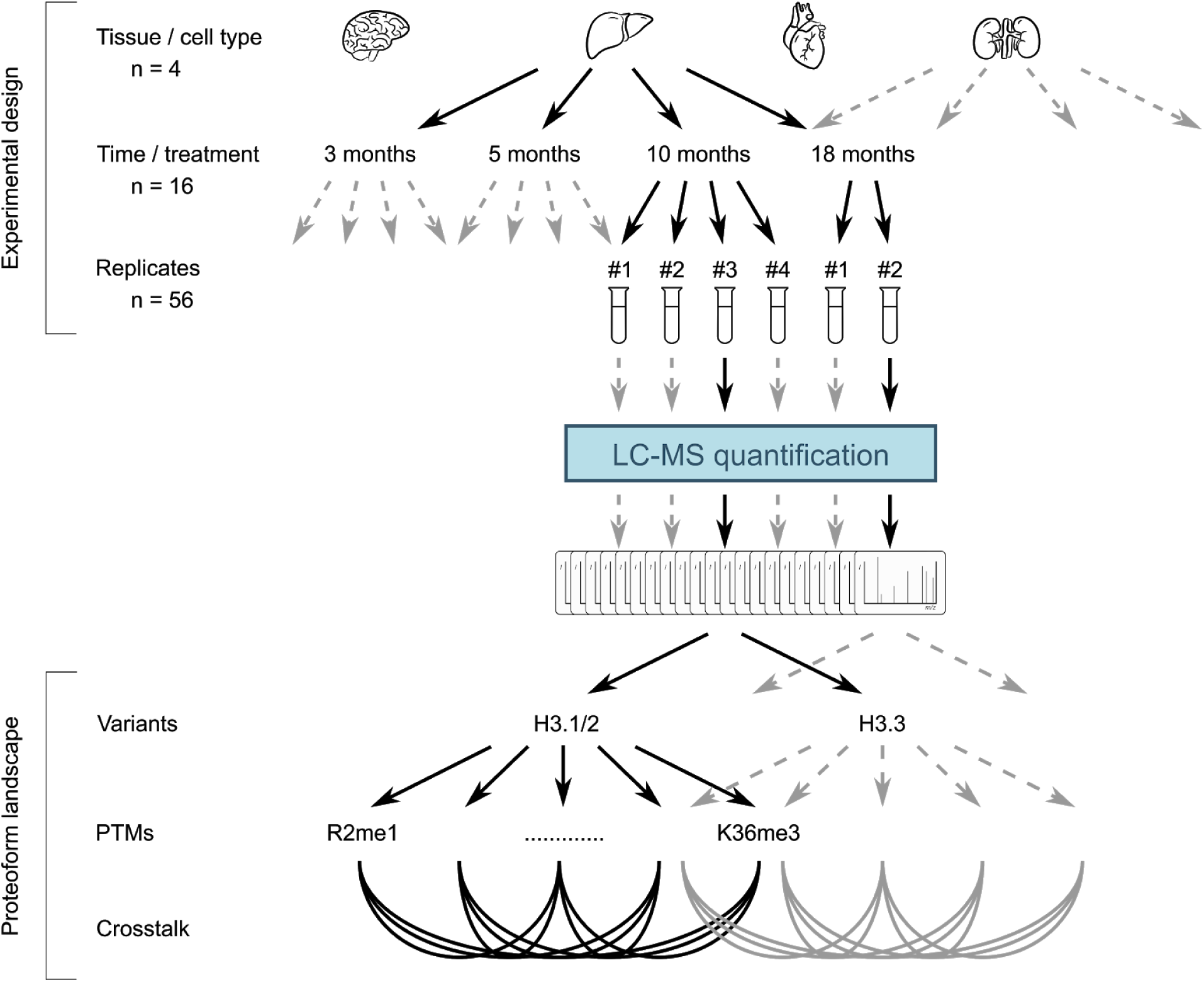
Complex experimental design of a proteomics study. This example investigated histone PTM dynamics and crosstalk in aging mouse tissues [9] but is representative in its complexity for many large-scale mass spectrometry studies nowadays. Especially the comparison of histone PTM crosstalk changes between different conditions and over time leads to very complex, multi-layered data that require novel visualization tools. Tissue icons by Saeful Muslim from thenounproject.com.

Specifically, we focus on the comparison of crosstalk dynamics of mono-, di- and trimethylated K9 and K27 of H3 in the four tissues (see Fig. 5). For simplicity, one separate PTM crosstalk map for each tissue was created, each containing the trajectories of all eight observed H3 PTM combinations from mice with ages from 3 months to 18 months. The PTM pairs are spread over the crosstalk landscape - four pairs show positive crosstalk in all tissues, the other PTMs are rarely observed together and characterized by strong negative crosstalk between for instance di- and tri-methylations. This confirms the highly diverse crosstalk patterns for different methylations states shown previously [10]. Note that the figure does not show the interplay between K9me3 and K27me3 as these two PTMs were not found to occur on the same histone and thus are mutually exclusive. H3K27 PTM abundances, as indicated by color code, vary substantially between methylation states, whereas the changes over time generally happen on smaller scale (as an example, also compare the line plot for K27me1 in heart in Fig. 3A). Intriguing is the strong change over time in the crosstalk between K9me1 and K27me1 (underlined) from strongly negative to neutral in the heart when compared to the other tissues. While the averaged negative interplay score over time, which was reported in the original heatmap [9], suggests a general negative crosstalk between these PTMs in heart, our crosstalk maps provide more detail. Following the development over time reveals a substantial change in crosstalk between K9 and K27 mono-methylation and that the two PTMs lose their mutually exclusive behavior with age. In addition, a comparison of K9me1K27me1 dynamics over different tissues shows that this PTM behavior is specifically found in heart, but not in brain, liver or kidney. In contrast, K9me2K27me3 (bold), which was identified as the relatively most abundant combinatorial PTM in 3 months old heart, kidney and liver in the original study [9], varies only slightly in both abundance and crosstalk, indicating a common role in stabilizing heterochromatin.

**Figure 5:**
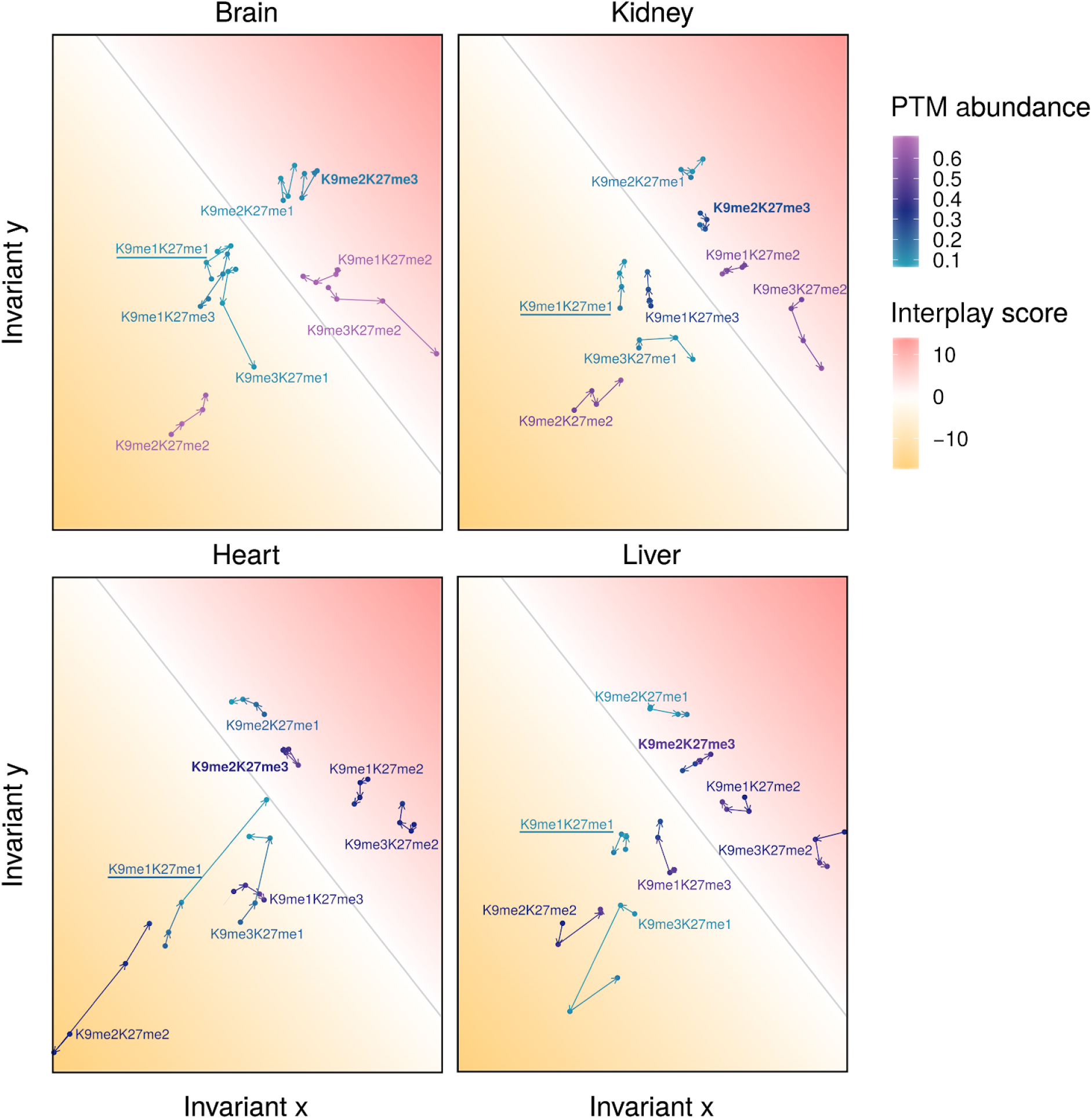
Visualization of PTM crosstalk dynamics in aging mouse tissues using PTM-CrossTalkMapper.

In conclusion, crosstalk maps circumvent summarizing PTM characteristics and thereby visualize PTM changes and differences in a more detailed manner than other PTM visualization approaches. At the same time, they provide a high-level overview over multi-dimensional PTM data and thereby new insights as well as directions for follow-up functional analyses to better understand complex epigenetic mechanisms.

The crosstalk maps allow an overall comparison of PTM abundances and crosstalk for all observed PTM combinations of methylated K9 and K27 positions between four tissues. Trajectories follow the PTM changes over tissue samples taken at 3, 5, 10 and 18 months of age. In this case, PTM abundance refers to the relative abundance of modifications at K27.

## Discussion

Middle-down mass spectrometry allows measuring histone proteoforms and their abundances with sufficient detail to quantify the crosstalk between distinct histone PTMs. There have been considerable efforts to visualize the resulting multi-layered, complex datasets that contain quantitative changes of tens of PTMs and their crosstalk, often compared over multiple experimental variables like tissue and age. These visualization approaches could focus on subsets of the information layers at a time or summarize the observed patterns by taking averages [10,12,13]. Specifically, crosstalk of multiple PTM pairs has previously been represented by hierarchical clustering of averaged interplay scores or simple networks, which did not allow showing crosstalk in direct context of the underlying PTM abundances and their dynamics [9,11].

We developed *PTM-CrossTalkMapper* to enable simultaneous visualization of PTM dynamics and PTM crosstalk across arbitrary sets of experimental conditions. PTM crosstalk maps can provide a high-level overview over the multiple layers of the PTM landscape and thereby valuable starting points and directions for further investigations of PTM crosstalk. At the same time, the maps provide greater detail than visualization approaches that require averaging over conditions, and thus provide insights into the functional roles of specific PTMs.

Given that proteins, especially histones, can carry multiple PTMs at the same time, the current focus of the interplay score on PTM pairs is a limiting factor in PTM crosstalk analysis. This can be solved by moving from visualizing the crosstalk between two PTMs to the crosstalk between multiply modified states. Alternatively, the interplay score can be extended to model crosstalk between more than two PTMs.

While the visualization framework was shown for histone PTM data, it is broadly applicable to any quantitative PTM dataset that includes absolute proteoform quantification. Going even further, the basic principle of defining a parameter landscape and data transformation could be adapted to facilitate the exploration of multi-dimensional, quantitative datasets in proteomics and other omics fields in general where crosstalk between subsets plays a biological role.

## Acknowledgements

R.K. was supported by a VILLUM Foundation postdoctoral fellowship (block-stipend) awarded to O.N.J. The O.N.J. laboratory at SDU is supported by generous grants to the VILLUM Center for Bioanalytical Sciences (VILLUM Foundation grant no. 7292) and PRO-MS: Danish National Mass Spectrometry Platform for Functional Proteomics (grant no. 5072-00007B).

## Notes

https://github.com/veitveit/CrossTalkMapper.

